# In depth amino acid mutational analysis of the key interspecific incompatibility transmembrane factor Stigmatic Privacy 1

**DOI:** 10.1101/2023.04.11.536390

**Authors:** Yoshinobu Kato, Shun Tadokoro, Shota Ishida, Maki Niidome, Yuka Kimura, Seiji Takayama, Sota Fujii

## Abstract

- In plants, there is an active prezygotic interspecific-incompatibility mechanism to prevent unfavorable hybrids between two species. We previously reported that an uncharacterized protein with four-transmembrane domains, named as Stigmatic Privacy 1 (SPRI1), is responsible for rejecting hetero-specific pollen grains in *Arabidopsis thaliana*.
- We have conducted a functional study of the SPRI1 protein, via point-mutational experiments and biochemical analysis. We studied the molecular regulatory mechanisms of SPRI1 and the relationships with its function.
- The alanine- and glycine-scanning experiments together with the evolutional analysis showed that the structural integrity of the C-terminal regions of the extracellular domain of this protein is important for its function. In addition, we found two cysteines (C67 and C80) within the extracellular domain that may be involved in the formation of intermolecular disulfide bonds. SPRI1 may form homo-multimers and is present as part of a ca. 300 kDa complex.
- Our present study indicates that molecular complex formation ability of SPRI1 may be important to maintain its stability and interspecific incompatibility functions in cells.

## Introduction

Pre-zygotic reproductive barrier is an important biological system that prevents formation of unfavorable interspecific hybrids (de Nettancourt, 2001). In plants, such reproductive barriers can be found in the process of inter-cellular communication between the male gametophyte pollen and the female pistil tissues (Tsuchimatsu & Fujii, 2022). Recently, we found a stigmatic protein called Stigmatic Privacy 1 (SPRI1) (Fujii *et al*., 2019), as the key factor for regulating hetero-specific pollen rejection in *Arabidopsis thaliana*. SPRI1 is a plasma-membrane localized protein with four-transmembrane domains. SPRI1 functions to reject pollen grains from other Brassicaceae species, and hetero-specific pollen tubes may grow into the mutants of SPRI1 while they are unable to do so in the wild-type stigma. It has been considered that pre-zygotic reproductive barrier can be broken down into incongruity and incompatibility. An incongruity arises due to the passive loss of gene functions involved in the fertilization process, probably acting as a driver of reproductive isolation during speciation. A pair of interactors, such as pistil cysteine-rich protein LURE1 and the cognate pollen-side receptor PRK6, has been known as the example of inter-specific incongruity causing factors in plant reproduction (Takeuchi & Higashiyama, 2016). On the other hand, SPRI1 is the first-reported interspecific incompatibility causing factor that actively rejects hetero-specific pollen (Fujii *et al*., 2019). More recently, a cysteine-rich peptide SpDIR1L in the *Solanum* species was found to be required to form interspecific reproductive barriers (Muñoz-Sanz *et al*., 2021). Thus, SPRI1 is one of the few known factors that form the pre-zygotic reproductive barrier through the incompatibility function. Therefore, there is great anticipation to uncover its cellular role.

Due to the lack of notable functional protein domains in SPRI1, its molecular role remains unclear. SPRI1 consists of four-transmembrane domains, two putative extracellular, and three intracellular regions. The four-transmembrane-based secondary structure of SPRI1 resembles that of the periplasmic protein DsbB in *E. coli* (Inaba & Ito, 2008). DsbB forms a disulfide bond in its periplasmic regions, and acts as a redox potential transducer across the cytoplasmic membrane. The tetraspanins are another protein family ubiquitously found in eukaryotic cells with secondary structures similar to SPRI1 or DsbB. Tetraspanins play a diverse role in cells such as regulation of signaling, cell-to-cell adhesion or fusion, fertilization and pathogen infection (Charrin *et al*., 2014). One of the proposed roles of the tetraspanins is to form a large molecular network in the membrane systems through direct or indirect protein-protein interactions, referred to as the tetraspanin ‘web’ in a previous study (Boucheix & Rubinstein, 2001).

In this study, we aimed to understand the biochemical function of SPRI1. In analogy with DsbB or tetraspanins, we suspected that the first 36 amino-acid-long extracellular regions in SPRI1 may play a critical role, and we performed a screening via point mutations. We show that disulfide bonds and complex formation abilities of SPRI1 are critical for its function to express interspecific incompatibility.

## Materials and Methods

### Plant materials and growth condition

All non-transgenic plant materials were obtained as describe in the previous study (Fujii *et al*., 2019). All plant materials were grown in mixed soil in a growth chamber under controlled conditions (light intensity 120–150 µmol m^-2^s^-1^, 14-h light/10-h dark cycle at 22 ± 2°C).

### Gene cloning, introduction of the mutations, deletions, and insertions

We used the *SPRI1A* genomic fragment carrying pCambia1300 binary plasmid (pCambia1300/*SPR1A* or pCambia1300/*SPRI1A-Venus*) created in our previous study (Fujii *et al*., 2019) as a template to introduce various mutations. We used a common QuikChange site-directed mutagenesis protocol mostly following the rationale in other studies (Liu & Naismith, 2008). In brief, 1 ng of pCambia1300/*SPR1A* was used as the template to perform the mutagenesis inverse PCR using KOD FX Neo polymerase kit (TOYOBO, Osaka, Japan). Inverse PCR in 50 µl reaction volume was performed with the following thermal routine: 98°C 30 sec denature, followed by 12 cycles of 98°C 10 sec; 55°C 5 sec; 68°C 10 min. Primers are listed in Table S1. The restriction enzyme *Dpn* I (Takara-Bio, Kusatsu, Japan) was added to the PCR product and the solution was incubated at 37°C for 1 hour, to eliminate the template plasmid. The solution was then used to transform NEB turbo *Escherichia coli* competent cells (NEB, MA, USA). pCambia1300/*SPRI1A-C+12* was created in our previous study (Fujii *et al*., 2019). The mutant plasmids were used to create the transgenic plants using the *Agrobacterium* infiltration procedure, as described in our previous study (Iwano *et al*., 2009).

### Pollination experiments

Flowers were emasculated before anthesis. At anthesis, pistils were harvested, placed on 1% agar plates, pollinated, and incubated for six hours at 22 ± 2°C, humidity 50±5%. Pistils were pollinated in a manner that entire stigmatic surface is covered with pollen grains. Pollinated pistils were fixed overnight at room temperature in ethanol/acetate 3:1 (v/v), then at 60°C for 30 minutes in 1M sodium hydroxide. The pistils were stained in 2% tripotassium phosphate/0.01% aniline blue for three hours at room temperature.

### Microscopic observation of pollen tubes

Pollen tubes on pistils stained with aniline blue were observed and counted under an epifluorescent microscope as previously described (Shiba *et al*., 2000). To facilitate large-scale phenotyping during the alanine scanning analysis, we defined arbitrary compatibility scores based on the numbers of pollen tubes in the styles: 1: No tubes observed; 2: 1–10 tubes; 3: 11–30 tubes; 4: 31–60 tubes; 5: ≥60 tubes.

### Microscopic observation of fluorescent proteins

Fluorescence in the emission range 520–555 nm was observed using a LSM880 confocal laser scanning microscope (Carl Zeiss, Oberkochen, Germany), with 514-nm excitation from an Argon laser.

### Procedures of obtaining antibodies against soluble parts of SPRI1

A synthetic gene encoding SPRI1 of which transmembrane domains (39th–61st, 100th– 122nd, 134th–156th, and 171st–193rd amino acids) were substituted by GGGGS linkers was purchased from FASMAC (Atsugi, Japan). This gene was amplified by PCR and inserted via In-Fusion system (TOYOBO) into the *Nde*I and *Xho*I sites of pET22b(+) (Novagen). Primers are listed in Table S1. Expression of the recombinant proteins was induced by 1 mM isopropyl β-D-thiogalactopyranoside at 37 °;C for 5 h in host *Escherichia coli* strain Rosetta (DE3) pLysS cells (Novagen). After induction, the cells were harvested in 20 mM potassium phosphate buffer (pH 8.0) containing 500 mM NaCl and cOmplete EDTA-free protease inhibitor cocktail (Roche). The inclusion bodies were pelleted from sonicated cells at 3000 × g for 15 minutes and solubilized in 20 mM potassium phosphate buffer (pH 8.0) containing 500 mM NaCl and 6 M guanidine hydrochloride. Insoluble material was removed by centrifugation at 10,000 × g for 1 h. The supernatant was incubated with COSMOGEL His-Accept (Nacalai, Kyoto, Japan) for 1 h. The beads were washed with 20 mM potassium phosphate buffer (pH 7.4) containing 500 mM NaCl and 4 M urea. The recombinant proteins were eluted with 20 mM potassium phosphate buffer (pH 7.4) containing 500 mM imidazole, 500 mM NaCl, and 4 M urea. Polyclonal antisera were raised against the purified recombinant protein in a mouse (T. K. Craft, Maebashi, Japan).

### Immunoblot analysis

Stigmas were collected from the Arabidopsis at flower stages (13-14) in 50 mM HEPES-KOH (pH 7.4) containing 5 mM MgCl_2_, 2 mM MnCl_2_, 10 mM NaF, 10 mM β-glycerophosphate, cOmplete ULTRA EDTA-free protease inhibitor cocktail (Roche), and 0.0125% (w/v) Tween-20 and stored at -−0°;C until use. One hundred stigmas were homogenized using a polypropylene pestle by hand in a 1.5 mL tube containing 100 µL of this buffer on ice. Debris was removed by 100 × g for 2 minutes, and supernatants were centrifuged at 20,000 × g for 30 minutes. Supernatants were collected as soluble fractions and used for detection of Actin. Remaining pellets were washed twice by resuspending in the 50 mM HEPES-KOH (pH 7.4) containing 5 mM MgCl_2_, 2 mM MnCl_2_, 10 mM NaF, 10 mM β-glycerophosphate, cOmplete ULTRA EDTA-free protease inhibitor cocktail and centrifugation (20,000 × g, 30 minutes). The final pellet was resuspended in 15 µL of the same buffer and used for detection of SPRI1A as a membrane fraction. After addition of equal amount of SDS sample buffer (50 mM Tris-HCl (pH6.8) containing 2% SDS, and 10% (v/v) glycerol with or without 50 mM DTT), samples were incubated at 60°;C for 15 min. TGX FastCast Acrylamide kit (12%, Bio-Rad, CA, USA) was used for SDS-PAGE. After electrophoresis, proteins were blotted on the Immobilon-P PVDF 0.45 µm (MERCK, Darmstadt, Germany). PVDF membranes were blocked using the PVDF Blocking Reagent for Can Get Signal (TOYOBO). Primary and secondary antibodies were diluted in the TBS-T buffer (0.05 M Tris-HCl (pH 7.6), 0.15 M NaCl, and 0.05% (v/v) Tween-20). Polyclonal antibody against Actin and GFP were purchased from Agrisera (AS13 2640, Vännäs, Sweden) and MBL (598, Tokyo, Japan), respectively, and used at a dilution of 1:5,000. Polyclonal antibody against SPRI1 was used at a dilution of 1:2,000. Secondary antibody against rabbit IgG (Goat Anti-Rabbit IgG (H+L)-HRP Conjugate (170-6515, Bio-Rad)) was used at a dilution of 1;25,000 for detection of Actin and Venus. Secondary antibody against mouse IgG (Goat Anti-Mouse IgG+IgM HRP conjugate (AMI0704, Biosource international, CA, USA)) was used at a dilution of 1;25,000 for detection of SPRI1.

### 2D BN/SDS-PAGE

Large pore BN-PAGE was performed as previously described (Järvi *et al*., 2011; Otani *et al*., 2018) with minor modifications. The pellet of membrane fractions prepared as described above were resuspend in 25 mM BisTris-HCl (pH 7.0) and centrifuged at 20,000 × g for 30 minutes. The pellet derived from two hundred stigmas was resuspended in 10 µL of 25 mM BisTris-HCl containing 20% (w/v) glycerol and 1% (w/v) *n*-dodecyl-β-D-maltoside for 5 minutes on ice. Insoluble materials were removed by centrifugation at 20,000 × g for 2 minutes. Supernatants were mixed with one-tenth volume of 100 mM BisTris-HCl (pH 7.0) containing 500 mM 6-aminocaproic acid, 30% sucrose, and 5% SERVA Blue G, and membrane protein complexes equivalent to two hundred stigmas were separated by 3% (T) 20% (C) acrylamide stacking gel and 3.5 – 12% (T) 3% (C) acrylamide gradient separating gel containing 50 mM BisTris-HCl (pH 7.0) and 500 mM 6-aminocaproic acid. NativeMark Unstained Protein Standard (Thermo Fischer Scientific, Massachusetts, USA) was also loaded. BN-gel was excised and incubated with SDS sample buffer (50 mM Tris-HCl (pH6.8) containing 2% SDS, and 10% (v/v) glycerol with or without 50 mM DTT) for 15 minutes at 60°;C. Gel strips were layered on TGX FastCast 12% Acrylamide gel. After electrophoresis, SPRI1 and Venus were detected as described above.

### Sequence analysis and dN/dS calculation of *SPRI1* by PAML

TMHMM server v2.0 (Krogh *et al*., 2001) was used to predict the transmembrane region of SPRI1. Sequences of the *SPRI1* orthologous genes were obtained as described in our previous study (Fujii *et al*., 2019). Multiple alignment of the SPRI1 proteins were performed by muscle (Edgar, 2004), followed by manual corrections. A codon-wise alignment of the *SPRI1* orthologs was performed using the tranalign function in the EMBOSS package. We used the site model in the Phylogenetic Analysis Using Maximum Likelihood (PAML) package to detect positively selected sites in the *SPRI1* orthologs. The dN/dS ratio was calculated with the two codon substitution models M7 and M8 included in the program codeml(Yang, 2007). M7 is a neutral model with continuous beta distribution of dN/dS values (≤1) is assumed. M8 overlaps with M7, except that it allows dN/dS > 1. M7 can be used as the null hypothesis against M8. All of the parameters in the codeml control file was default with ‘model’ set as 0. The fitness of codon substitution models was evaluated with log-likelihood ratio statistics, which are assumed to be χ^2^ distributed. Bayes Empirical Bayes (BEB) probabilities of positive selection and dN/dS values for each site were found from the rst output file from the codeml run.

### The statistical analysis

All of the statistical analysis was done using R (R Core Team, 2017). For all box plots, center lines show the medians, box limits indicate the 25th and 75th percentiles, whiskers extend 1.5 times the interquartile range from the 25th and 75th percentiles, and data points are plotted as open circles.

## Results

### Amino acid substitutions in the long extracellular domain of SPRI1 highlights amino acids required for its stability

The SPRI1 protein contain two putative extracellular domains based on its anticipated membrane insertion topology (Figure S1). We paid attention to the longer domain, which is predicted to span from the 63rd glutamine (Q63, amino acid residues are abbreviated in this manner hereafter) to F98 (Figure 1a). To understand the role for each of these residues, we substituted 32 of these amino acid residues by alanine and introduced these mutant SPRI1A (functional allele of SPRI1 found in the previous study) forms into the *spri1-1* mutant. The lines expressing the alanine-replaced SPRI1A forms were pollinated with the pollen grains of *Malcolmia littorea*, a species belonging to the Brassicaceae family whose pollen has been shown to be rejected by the function of SPRI1 in our previous study (Fujii *et al*., 2019). As a result, we found that replacements of amino acids C67, C80, G85, T86, I88, H91, K95 and R96 have caused significant impairment of the function of SPRI1 *in vivo* (Figure 1b).

**Figure 1.**
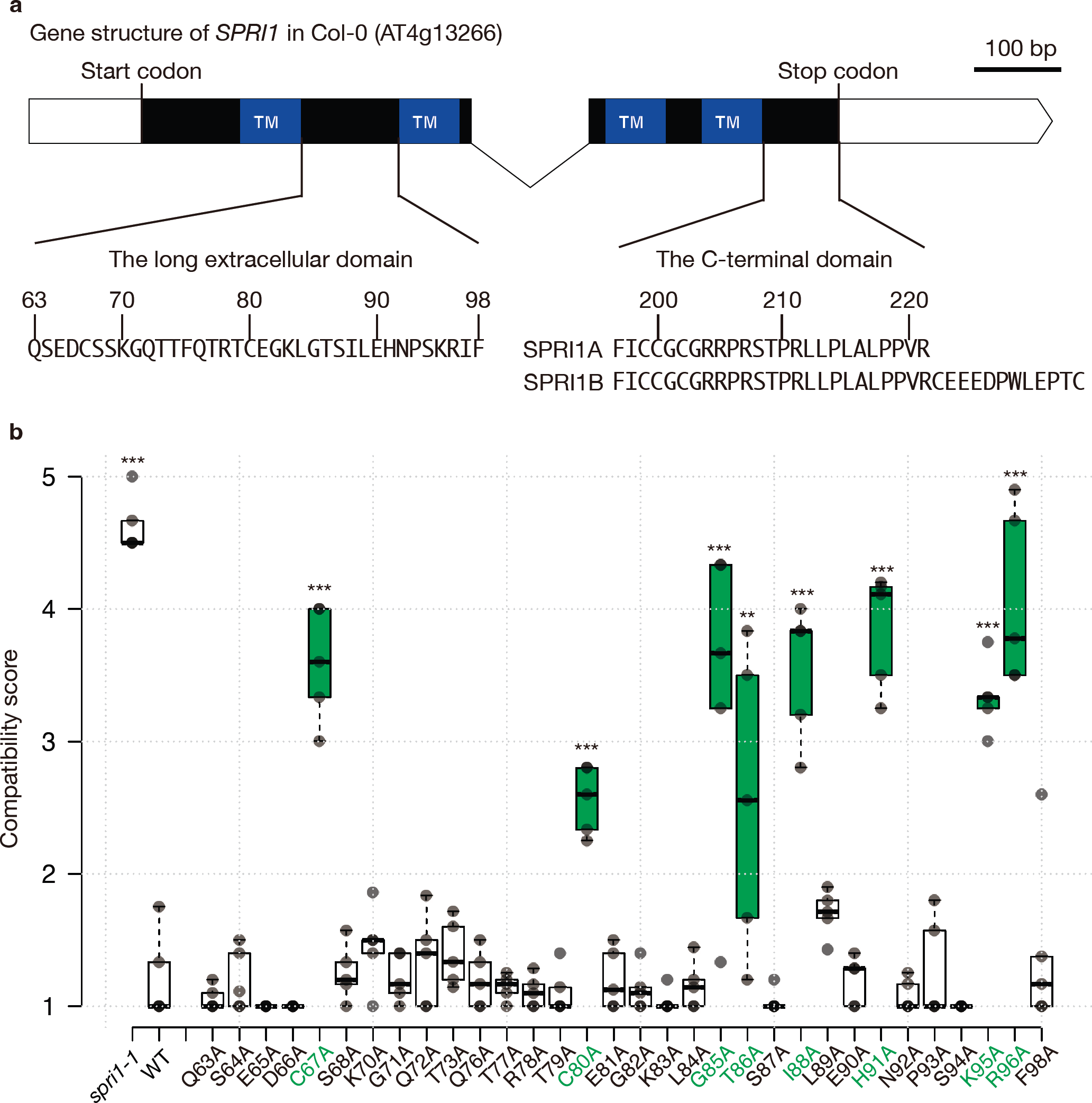
Gene structure of SPRI1 and alanine replacement analysis of the long extracellular domain. **(a**) Gene structure of SPRI1. TM: transmembrane region. (**b)** Summary of the interspecific pollination analysis of the transformants introduced with the alanine-replacement mutant SPRI1 forms. Significant differences by Dunnett’s test compared against wild-type (WT) are indicated by **(*p* < 0.01) and ***(*p* < 0.005).

### Structural stability at the evolutionary conserved region in the long extracellular domain is important for SPRI1 function

From the above result, we noticed that replacements of the C-terminal half of the long extracellular domain (arbitrarily named as the LEDC-region, Figure 1a) with alanine cause frequent impairment to the SPRI1 function. We reasoned that this could be due to a stronger evolutionary pressure to preserve the protein structure on the LEDC-region compared to the N-terminal region of the long extracellular domain (LEDN-region). Thus, we analyzed the sequences from 58 *SPRI1* orthologous genes collected in our previous study (Fujii *et al*., 2019), and calculated the ratio of non-synonymous to synonymous substitution rates (dN/dS) (Figure 2a). We observed that the dN/dS ratio for amino acid positions in the LEDN region frequently exceeded 1, indicating that these codons are either experiencing relaxed selective pressure or are under positive selection (Figure 1a). We were unable to obtain dN/dS ratio for some residues due to amino acid insertion/deletions in this region, further implying that this LEDN-region could be diverged. In contrast, dN/dS ratio of most codons in the LEDC-region are found to be <1 (Figure 2a), suggesting that they are more likely to be under negative selection and are structurally conserved.

**Figure 2.**
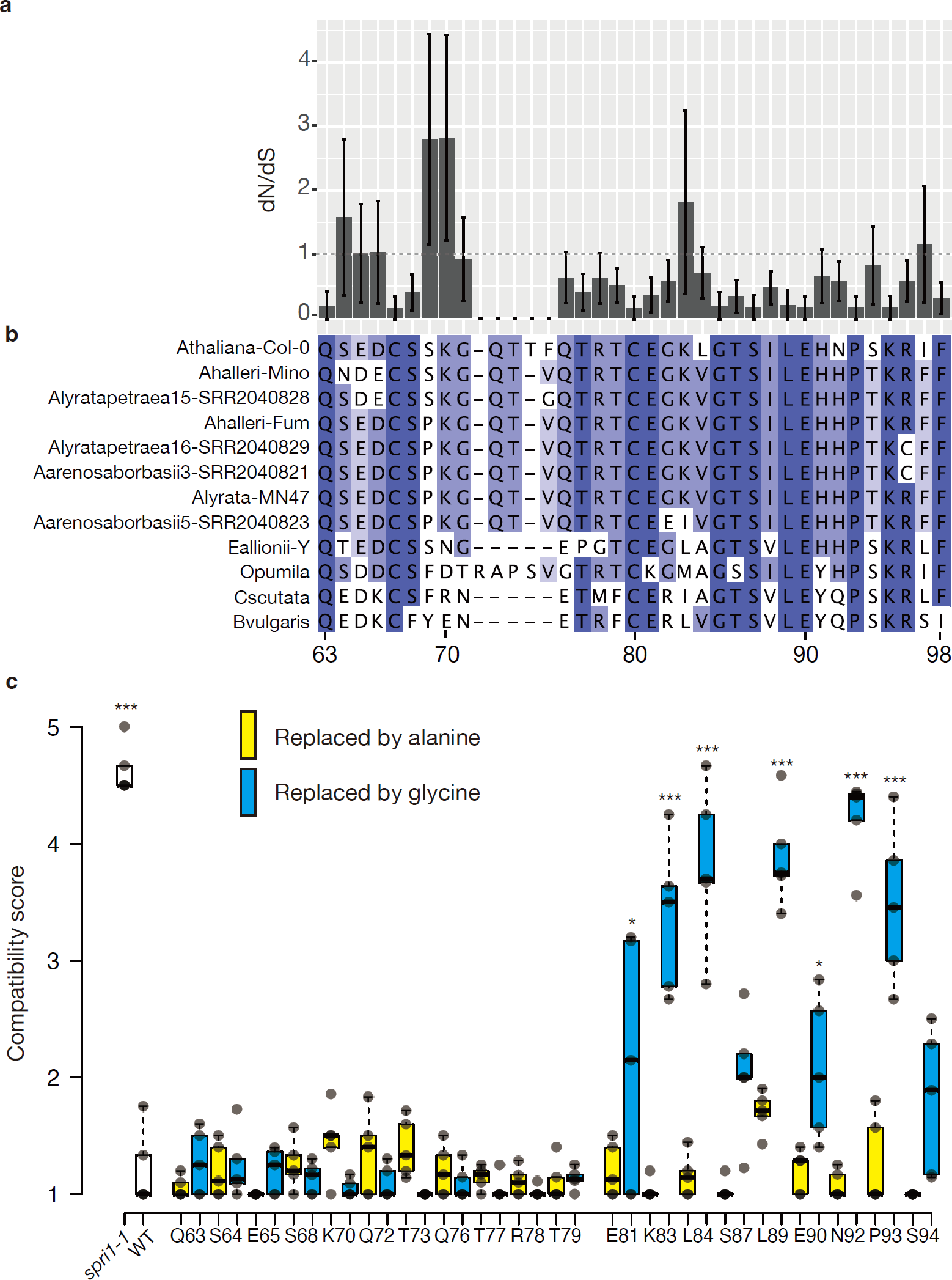
Evolutionary analysis of SPRI1 and glycine replacement analysis of the long extracellular domain. **(a)** Bargraph showing dN/dS calculated from the SPRI1 multiple alignment from 58 Brassicaceae species. **(b)** Amino acid sequence alignment of the long extracellular domain found from representative Brassicaceae species. (**c**) Summary of the interspecific pollination analysis of the transformants introduced with the alanine-replacement or the glycine-replacement mutant SPRI1 forms. Significant differences by Dunnett’s test compared against wild-type (WT) are indicated by *(*p* < 0.05) and ***(*p* < 0.005).

Next, we asked if increasing protein structural flexibilities in these regions would indeed impair the SPRI1 function. We selectively replaced the amino acid positions that were able to be replaced by alanine without any functional compromises (Figure 1b) with glycine, the smallest amino acid which renders largest conformational flexibility to a protein structure. As a result, while replacement with glycine in the LEDN-region did not exhibit any effect, most of the glycine replacements in the LEDC-region impair the molecular function of SPRI1A (Figure 2c). Taken together with the evolutionary analysis, these data indicated that the C-terminal region of the extracellular domain are structurally important to maintain the function of SPRI1.

### Two cysteine residues in the long extracellular domain may be involved in forming the disulfide bonds

We also found that the two cysteine residues C67 and C80 were the only indispensable amino acids in the LEDN-region (Figure 1b). We suspected that these residues could be involved in intra- or inter-molecular disulfide bond formation. To investigate the possible involvement of disulfide bond formation in SPRI1, we performed immunoblots in the absence (-) or the presence of the reducing agent dithiothreitol (DTT). When treated with DTT, a single band was detected close to its expected monomeric molecular size 25.1 kDa (Figure 3a). In contrast, in the absence of DTT, we detected an additional band close to the 37 kDa marker (indicated by the arrow) and a faint band above the 50 kDa marker (indicated by the arrowhead) (Figure 3a). It was possible that these bands correspond to homo-polymers, and SPRI1 may form inter-molecular disulfide bonds with itself. To verify this hypothesis, we developed a transgenic *spri1-1* line expressing the fusion protein of SPRI1A and a fluorescent protein Venus (Nagai *et al*., 2002). We reasoned that if SPRI1–SPRI1 disulfide bond is formed, the size shift of SPRI1A–Venus in the non-reducing condition will become greater because of the Venus tag fusion. We confirmed that this SPRI1A–Venus line can strongly reject pollen grains of *M. littorea* (Figure S2), indicating that the native function of SPRI1A is retained in this transgenic line. In both presence and absence of DTT, the SPRI1A–Venus signals detected by the anti-GFP antibody were found at around its expected monomeric molecular size 52.7 kDa (Figure 3b). In the absence of DTT, an additional band at around >100 kDa was strongly detected (Figure 3b). This size shift observed in the SPRI1A–Venus transgenic line (Figure 3b: about 50 kDa) was greater than the native SPRI1A (Figure 3a: about 10–15 kDa) supported the idea that the higher molecular band (indicated by the arrow) in the non-reducing condition may be composed of SPRI1A-homo-dimer. The apparent molecular size shift in the native SPRI1A dimer was smaller than its expected size (25 kDa + 25 kDa), probably due to the incomplete denaturation and compact packing under DTT-less condition.

**Figure 3.**
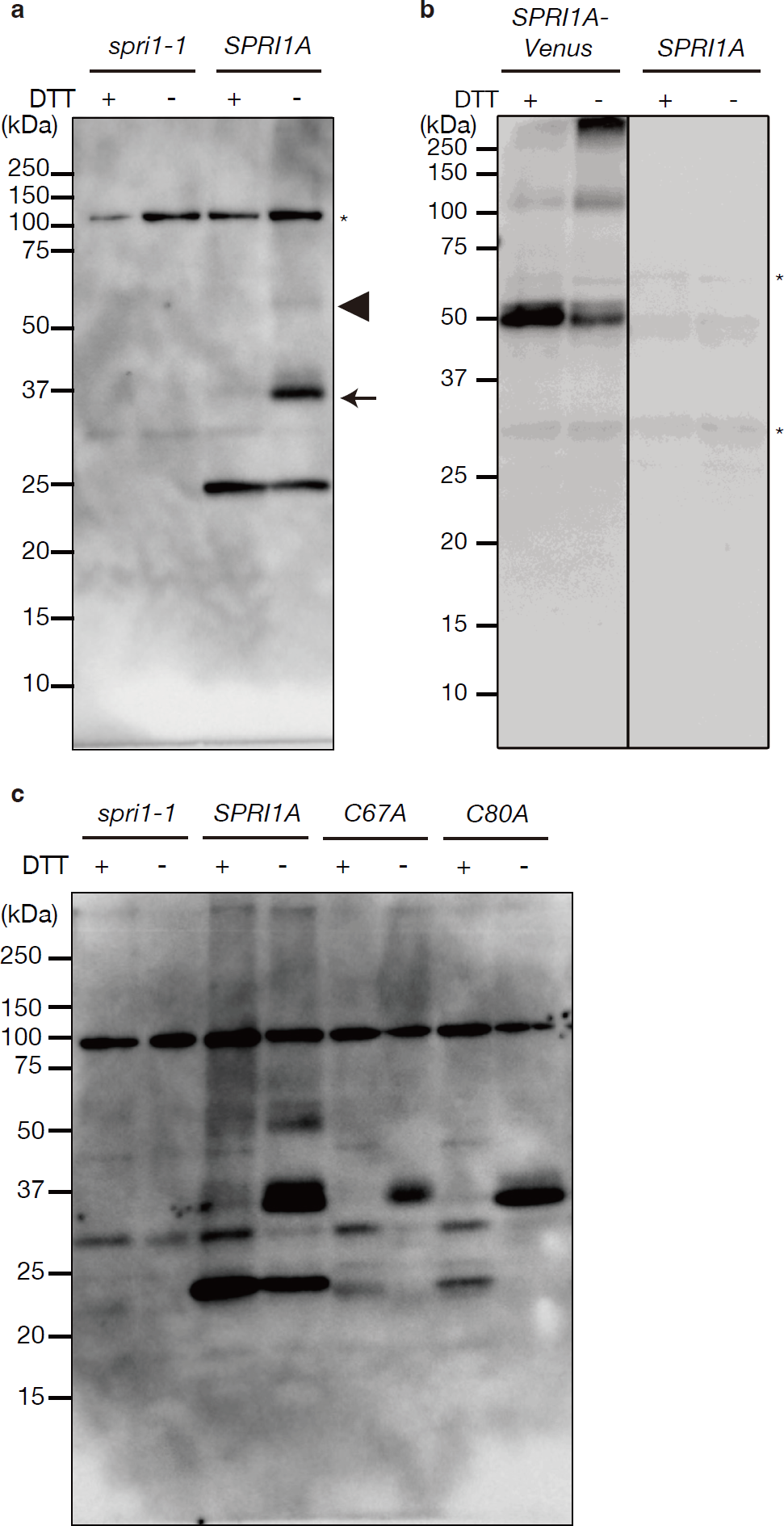
SPRI1A forms disulfide bonds. **(a)** Total membrane proteins were separated under the presence (+) or absence (-) of DTT conditions in SDS-PAGE followed by immunoblotting using the antibody against SPRI1. *spri1-1* was used as a negative control. A black arrow and an arrowhead indicate the putative SPRI1 dimer and some higher order structure, respectively. A black asterisk indicates non-specific signal. **(b and c)** The same analysis as in Figure 3a was performed using *SPRI1A– Venus* with the antibody against GFP in panel b and using *SPRI1A_C67A* (*C67A*) and *SPRI1A_C80A* (*C80A*) with the antibody against SPRI1 in panel c. The uncropped image of panel b was shown in Figure S5.

To further investigate the involvement of C67 and C80 in the disulfide bond formations, we detected the molecular accumulation pattern of SPRI1A protein in the alanine-substitution lines of these two amino acids. Unlike other amino acid mutations (H91A, K95A, R96A) that affected the accumulation of SPRIA itself, lines expressing C67A and C80A accumulated significant amount of SPRI1 protein (Figure 3c, Figure S3). In non-reducing condition, only the putative dimer forms were detected in the C67A and C80A mutants (Figure 3c). This may indicate that both C67-C67 and C80-C80 disulfide bonds could be formed. The slight mobility difference of the putative dimeric bands detected in the two mutants (Figure 3c, Figure S3) may reflect the difference of the molecular conformations in the C67-C67 (detected in C80A) and C80-C80 (detected in C67A) forms. These results suggested that SPRI1A may form inter-molecular disulfide bonds.

On the other hand, SPRI1A monomer was not detectable under absence of DTT in these mutants, although it was stably accumulated in the plant expressing native SPRI1A (Figure 3c). It is possible that an intra-molecular disulfide bond between C67 and C80 is also formed and contributes to the SPRI1 stability. To test this possibility, we produced a transgenic plant expressing SPRI1A of which both C67 and C80 were substituted to alanine and detected the SPRI1A protein in the transformants. In this transgenic plant, the accumulation level of SPRI1A was drastically reduced to less than 1/8 compared to the wild-type SPRI1A (Figure 4a). As opposed to the single point-mutated C67A or C87A plants, only the monomeric SPRI1A form was detected in the C67A_C80A plant under the non-reducing condition (Figure 4b). Since this monomeric SPRI1A can stably accumulate in the wild-type plant, it is possible that intra-molecular disulfide bond between C67 and C80 make SPRI1A stable. Taken together, both intra- and inter-molecular disulfide bonds are formed via C67 and C80, and they are crucial for stabilizing SPRI1.

**Figure 4.**
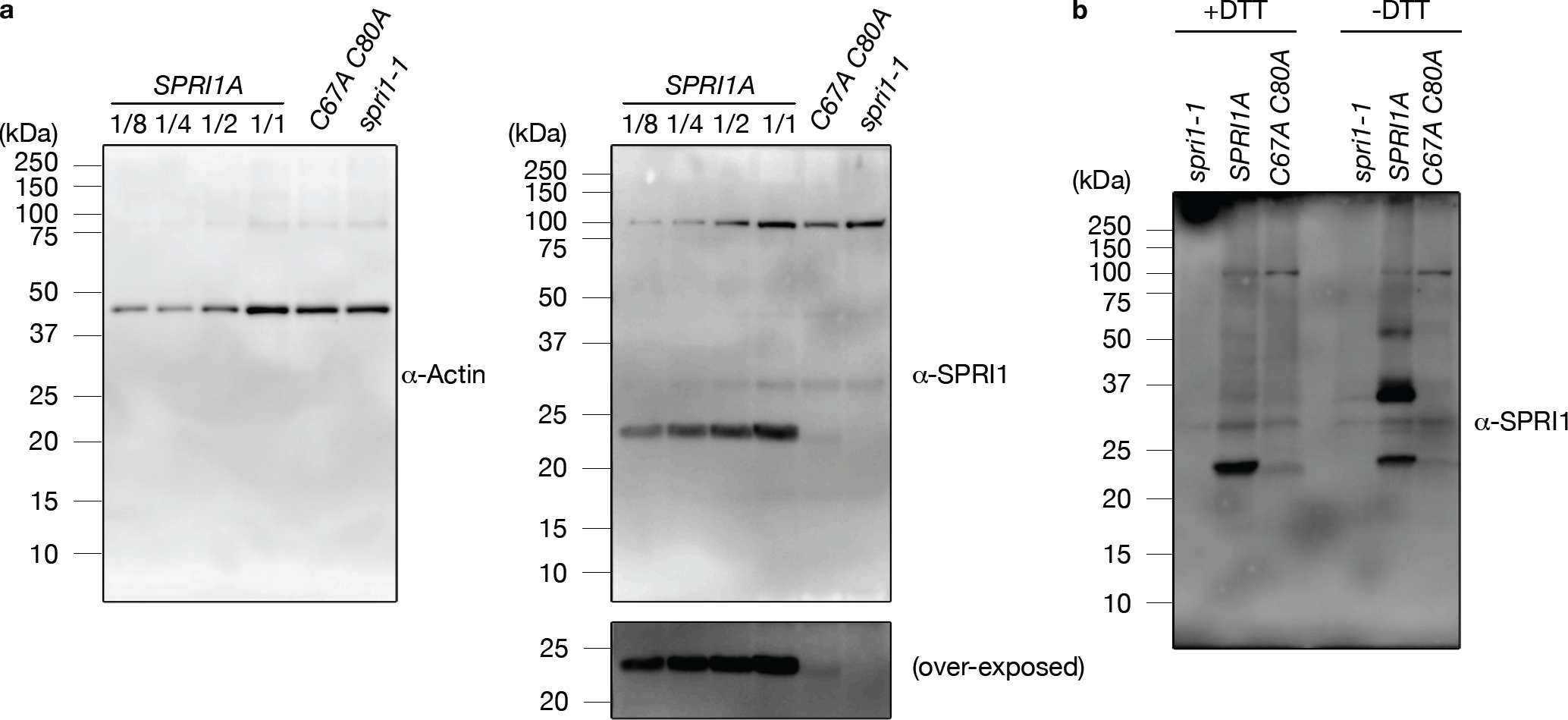
C67 and C80 are essential for the SPRI1 accumulation. **(a)** Total membrane proteins isolated from stigma samples of *SPRI1A*, *SPRI1A_C67A C80A* (*C67A C80A*), and *spri1-1* were analyzed by immunoblotting using antibody against SPRI1. Sample loading was based on the number of stigmas, along with a dilution series of *SPRII1A*. Total soluble proteins were also analyzed, and actin was detected as a loading control using its specific antibody. **(b)** The same analysis as in Figure 3a was performed using stigma samples of *spri1-1*, *SPRI1A*, and *SPRI1A_C67A C80A* plants.

### SPRI1 is found within a high-molecular complex including its homo-multimer forms

We next suspected that SPRI1 could reside in a higher-order complex. We therefore used the Blue Native-PAGE (BN-PAGE) to study the native SPRI1 protein complex, and the possible recruitment of SPRI1 homo-polymers therein. We used the mild detergent n-dodecyl-β-D-maltoside to solubilize the SPRI1 complex from the membranes in the homogenized stigmatic cells and subjected to the two-dimensional BN/SDS-PAGE. As a result, SPRI1 was detected at the size between 242 kDa and 480 kDa in the BN-gel (Figure 5a). These signals were not detected in the *spri1-1* mutant (Figure 5b), indicating that they are specific signals of SPRI1A. The SPRI1 monomer and the putative homo-dimers were detected even in the DTT treated BN-gel strip, probably because enough amount of DTT could not reach the SPRI1 proteins in the gel strip during the SDS sample buffer treatment. In contrast, we observed SPRI1 signals near 720 kDa in the SPRI1–Venus expressing line under the same BN-PAGE conditions (Figure 5c). This finding suggests that the putative SPRI1 complex increased in size substantially by fusing Venus to SPRI1 monomers. This result implies that the high-order SPRI1-multimer likely constitute the primary component of this complex (Figure S4a), rather than the possibility that proteins other than SPRI1 occupy majority this complex (Figure S4b).

**Figure 5.**
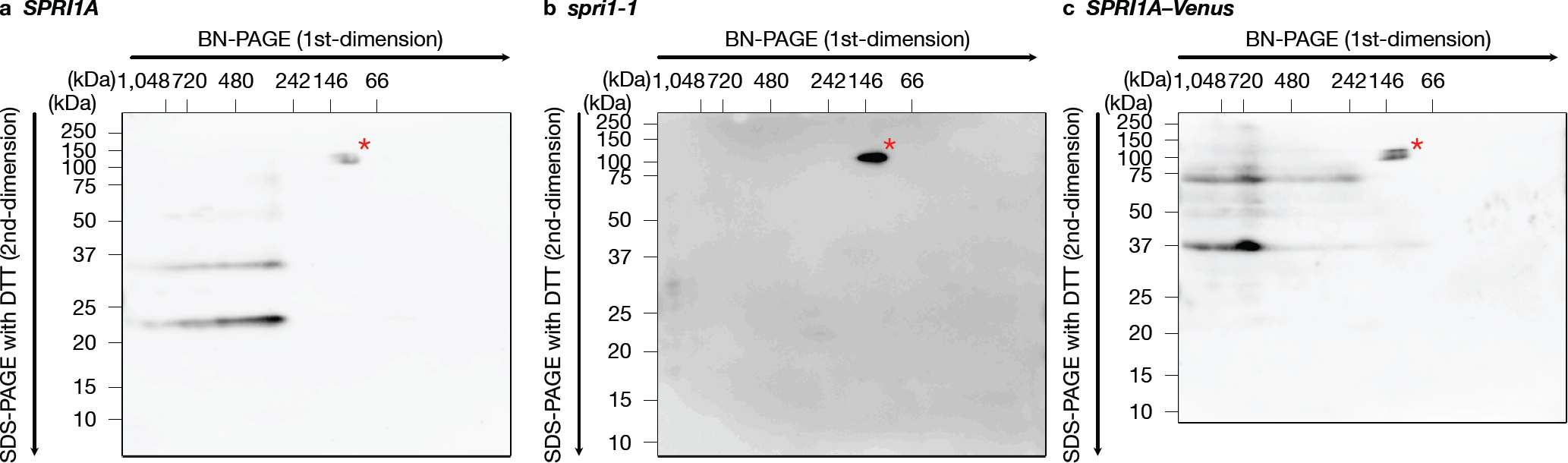
SPRI1A forms a protein complex of which size is about 300 kDa. **(a-c)** Stigmatic membrane protein complexes derived from *SPRI1A* (a), *spri1-1* (b), and *SPRI1A–Venus* (c) were separated by BN-PAGE and further subjected to two-dimensional SDS-PAGE under the presence of DTT condition. The SPRI1 protein was detected using the antibody against SPRI1. Red asterisks indicate non-specific signals. It was observed that the migration of the SPRI1A-Venus monomer and putative SPRI1A-Venus dimer in two-dimensional SDS-PAGE was faster than in standard SDS-PAGE (Figure 3b). This was probably due to the Venus moiety being undenatured without TCA treatment, which cannot be applied to a BN-gel strip.

In addition, SPRI1 was detected in the absence of DTT in two-dimensional SDS-PAGE (Figure 6). The putative SPRI1 protein complex (approximately 300 kDa) exhibited various redox states of SPRI1A, including intra- and inter-molecular disulfide bonds (Figure 6a). This observation suggests that the SPRI1 homo-oligomer is a mixture of SPRI1 monomers, homo-dimers, and higher molecular structures linked by disulfide bonds. The putative SPRI1 protein complex (ca. 300 kDa) was also present in the C67A and C80A mutants (Figure 6b and 6c), but it consisted purely of SPRI1 inter-molecular dimers. This indicates that the formation of SPRI1-multimers is not dependent on specific disulfide bond structures. It is consistent with the idea that the SPRI1-multimer is a mixture of various redox states of SPRI1.

**Figure 6.**
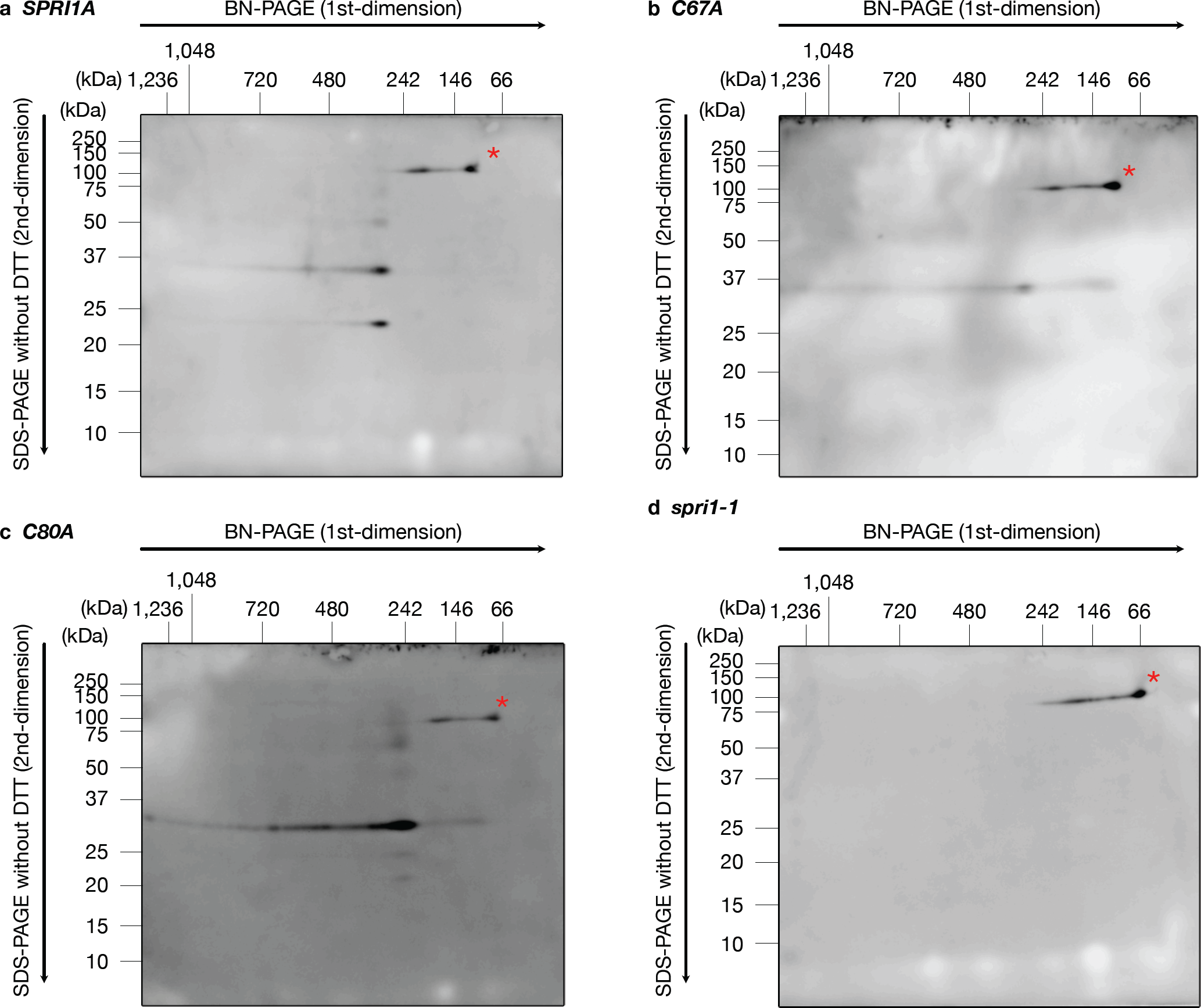
The SPRI1 protein complex contains both intra- and inter-molecular linked SPRI1. **(a-d)** The almost same analysis as in Figure 5 was performed. Two-dimensional SDS-PAGE was conducted under the absence of DTT condition, and the SPRI1 protein was detected using the antibody against SPRI1. Stigma samples of *SPRI1A* (a), *SPRI1A_C67A* (b), *SPRI1A_C80A* (c), and *spri1-1* plants were used.

### C-terminal sequences and putative palmitoylation sites are also required for the function of SPRI1

Apart from the long extracellular domain, we also reinvestigated the function of the C-terminal domain in this study. In our previous works, we found that some *Arabidopsis* natural strains (haplotype *SPRI1B* carriers) had lost their SPRI1 function due to the addition extra 12-amino acids at the C-terminus caused by a frameshift mutation (Fujii *et al*., 2019) (Figure 7a). We showed that addition of this C-terminus extension to the functional SPRI1A can compromise its function, leading to the idea that this region is functionally important. To further investigate the regulation of SPRI1A by the C-terminal region, we created a series of deletion and extension lines (Figure 7a) and introduced these mutant SPRI1A forms into the *spri1-1* mutant. We found that deletion of 9 amino acids, or extension of 7 amino acids at the C-terminus of SPRI1 caused severe defects to its function (Figure 7b). In some of the lines (C+7, C+9, C+11, C+12), we found that SPRI1 protein accumulation level was below the detection limit, although SPRI1A fused with Venus at its C-terminus can fully complement its function accumulation (Figure S2a). Thus, it was possible that some specific amino acid sequence in the C-terminal extension can cause protein instability (Figure 7c).

**Figure 7.**
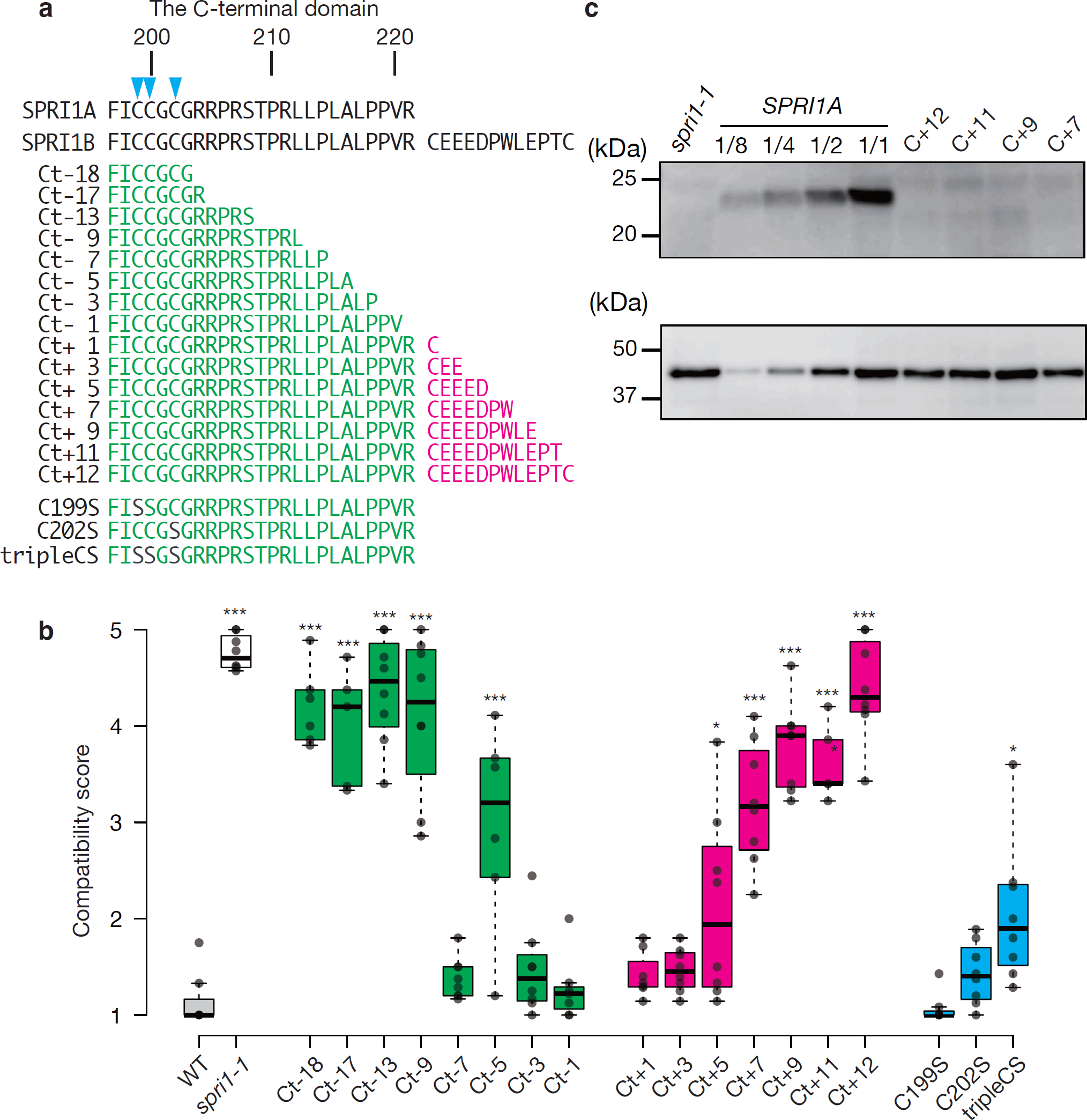
The intact C-terminal sequence is essential for the SPRI1 function. **(a)** Diagram showing the C-terminal region of SPRI1 and the deletion or the extension mutant series. Blue arrow heads indicate the cysteine residues which were predicted as palmitoylation sites. **(b)** Summary of the interspecific pollination analysis of the transformants introduced with the C-terminal deletion series (C18 to C-1), the extension series (C+1 to C+12) or the putative palmitoylation sites. Significant differences by Dunnett’s test compared against wild-type (WT) are indicated by *(*p* < 0.05) and ***(*p* < 0.005). **(c)** Total membrane proteins isolated from stigma samples of *SPRI1A*, spri1-1, and a series of C-terminal extension lines (C+12, C+11, C+9, and C+7) were analyzed by immunoblotting using antibody against SPRI1. Sample loading was based on the number of stigmas, along with a dilution series of *SPRII1A*. Using total soluble proteins, actin was detected as a loading control with its specific antibody.

In tetraspanin proteins such as CD9, post-translational palmitoylation of the juxtamembrane cysteine residues plays an important role in the protein-protein interactions (Charrin *et al*., 2002). Since juxtamembrane cysteine residues at the 199, 200 and 202 positions of SPRI1A were strongly predicted as the candidate palmitoylation sites (Figure 7a) using CSS-Palm 4.0 (Ren *et al*., 2008), we replaced these amino acids with serine to investigate their contributions to the SPRI1A function. As a result, although single replacements (C199S, C202S) did not significantly affect the SPRI1A function, mutant of all three sites was partially but significantly impaired in the SPRI1A (Figure 7b). These results suggested that the C-terminal intracellular flexible region of SPRI1A is important for its function, as well as the putative palmitoylation sites.

## Discussion

In this study, we used mutant analysis to investigate the molecular nature of the SPRI1 protein. In the long extracellular region, we found that there is an evolutionary conserved subregion (LEDC-region) required for the function of SPRI1 (Figures 1 and 2). We also found that length of the flexible C-terminus regions is also a critical factor for the interspecific pollen rejection ability of SPRI1 (Figure 7). These regions may be required to stabilize the SPRI1 molecule in vivo, because we failed to detect detectable amount of SPRI1 protein in many of the mutants in these regions (Figure 7c, S3).

We also discovered that two cysteine residues within the long extracellular region are crucial for SPRI1 function in the stigma (Figure 1). It was possible that cysteine residues participate in the formation of intra- and inter-molecular disulfide bonds (C67-C67 and C80-C80) of SPRI1 in the stigma (Figures 3 and 4). We found that the C67A mutation had a more severe effect on the interspecific incompatibility phenotype compared to the C80A mutation (Figure 1b). This may be correlated with the observation that C67-C67 bridged inter-molecular dimer form of SPRI1 may be relatively more stable than the C80-C80 mediated form (Figure S3), indicating that C67-C67 plays a more significant role in the stability of SPRI1 in cells. Our observation also suggests the formation of intra-molecular C67-C80 disulfide bonds, as the monomeric form was absent in the C67A and C80A mutants, while it was abundant in the wild-type (Figure 3c). Since impairment of interspecific incompatibility was observed in the C80A plant, where the total amount of SPRI1 was unchanged (Figures 1 and S3), it is likely that C67-C80 intra- and/or inter-molecular disulfide bond is required for the full function of SPRI1. Recently, crystal structure of the four-transmembrane protein CD9 was revealed (Umeda *et al*., 2019, 2020). CD9 was found to form two intra-molecular disulfide bonds via four well-conserved cysteine residues in its large extracellular region, and these bridges were considered as important to stabilize this region (Umeda *et al*., 2020). It is possible that similar intra-molecular disulfide bond mechanism stabilizes the SPRI1 extracellular region. Regarding tetraspanins, palmitoylation of the cysteines at the end of transmembrane helices of CD9 is considered as important to anchor this protein to the lipid bilayer (Umeda *et al*., 2020). Similar juxtamembrane cysteine residues of SPRI1 (C199, C200, C202) were found to be important for its full function (Figure 7b), suggesting the commonality in the regulatory mechanism of SPRI1 and tetraspanins.

Relocation of intra and inter molecular disulfide bonds of the four transmembrane protein DsbB in *E. Coli* has been reported and was proposed to regulate the function of this protein (Inaba *et al*., 2006). It is also possible that intra- and inter-molecular disulfide bond switching occur in SPRI1 and may regulate its function.

The protein complex of approximately 300 kDa is formed by SPRI1, consisting of a mixture of intra-molecular linked SPRI1 monomer and inter-molecular linked SPRI1 dimers, suggesting that it is a SPRI1 homo-oligomer. Notably, this complex was also observed in C67A and C80A plants where almost all SPRI1 was inter-molecular bridged (Figures 6b and 6c), indicating that the SPRI1-multimer contains an even number of SPRI1. One plausible function of this protein complex is acting as a molecular hub, where SPRI1 interacts with other proteins, such as tetraspanin proteins. (Boucheix & Rubinstein, 2001). The Brassicaceae self-incompatibility system triggers calcium signaling upon self-pollination (Iwano *et al*., 2015), which might also occur in rejecting hetero-specific pollen grains. Given that SPRI1 has the ability to discriminate the species origin of pollen (Fujii *et al*., 2019), it is reasonable to assume that SPRI1 modulates the activity of its interacting partner(s) upon perceiving pollen grains.

## Acknowledgements

We thank M. Ishii, K. Mori, M. Saito, and A Yoshida for their technical assistance. This work was supported in part by Grants-in-Aid for Transformative Research Areas (22H05174 to S.F.), Grants-in-Aid for Scientific Research on Innovative Areas (16H06467, 16H06464 to S.Tak.; 16H01467 to S.Fuj.), Grants-in-Aid for Scientific Research (16H06380, 21H04711, 21H05030 to S.Tak.; 18H02456 to S.Fuj.), Grant-in-Aid for Challenging Exploratory Research (15K14626 to S.Fuj.) from the Ministry of Education, Culture, Sports, Science and Technology of Japan (MEXT), research fellows (17J09745, 19J01563 to Y.Kat.; 20J01572 to Y.Kim.) from the Japan Society for the Promotion of Science, Japan Science and Technology Agency (JST) PRESTO program (JPMJPR16Q8 to S.Fuj.; JPMJPR21D4 to Y.Kat.), and the Suntory Rising Stars Encouragement Program in Life Science (to S.Fuj.). The authors declare no competing financial interests.

## Author contributions

S.Fuj. and S.Tak. conceived the study. Y. Kat, S.Tak., and S.Fuj. wrote the manuscript. S.Ish, M.Nii., Y.Kim., and S.Fuj. conducted pollination experiments including production of transgenic plants. S.Ish. and Y.Kim. conducted microscopic observation of fluorescent proteins. S.Fuj. conducted *in silico* sequence analysis and dN/dS calculation. Y.Kat. conducted raising the antibody against SPRI1. Y.Kat. and S.Tad. conducted the immunoblot analysis and BN-PAGE.

## Materials and correspondence

Sota Fujii

## Legends of the Supplementary Figures and Table

**Supplementary Figure 1.**

A prediction result of the transmembrane regions of SPRI1 using TMHMM

**Supplementary Figure 2.**

**(a)** The interspecific pollination assay in *spri1-1* and *spri1-1*/SPRI1A-Venus. Significant differences by Dunnett’s test compared against *spri1-1* are indicated by ***(*p* < 0.005). **(b)** Representative images of the interspecific pollination assay after aniline blue staining. **(c)** The expression of Venus-fused SPRI1A at stigma was confirmed at stigmatic papilla cells in *spri1-1*/SPRI1A-Venus. *spri1-1* was observed as a negative control.

**Supplementary Figure 3.**

Total membrane proteins isolated from stigma samples of *SPRI1A*, *spri1-1*, and the lines expressing the alanine-replaced SPRI1A (*H91A*, *K95A*, *R96A*, *C67A*, *C80A*) were analyzed by immunoblotting using antibody against SPRI1. Because of the destabilization of SPRI1A_C67A and SPRI1A_C80A after addition of the SDS sample buffer including DTT, these samples were analyzed in the non-reducing condition. Sample loading was based on the number of stigmas, along with a dilution series of SPRII1A. Using total soluble proteins, actin was detected as a loading control with its specific antibody.

**Supplementary Figure 4.**

Schematic of possible SPRI1 complex models. SPRI1 is indicated in blue and orange, while other unknown factors within the complex are indicated in brown. (a) The SPRI1 multimer is the major component in this model. Attachment of multiple Venus molecules to the SPRI1 multimer results in a significant enlargement of the SPRI1 complex, consistent with the observation in Fig. 5, where a large difference in molecular complex size was found between SPRI1 and SPRI1-Venus. (b) The major component(s) of the SPRI1 complex in this model is the unknown factor(s) that interacts with SPRI1. However, attachment of only a few Venus molecules causes only a slight increase in the SPRI1 complex, which disagrees with the observation in Fig. 5.

**Supplementary Figure 5.**

The uncropped image of Figure 3b.

**Supplementary Table 1.**

The list of primers used in this study

