## Supplementary Figures for "In depth amino acid mutational analysis of the key interspecific incompatibility transmembrane factor Stigmatic Privacy 1"

**Supplementary Figure S1**

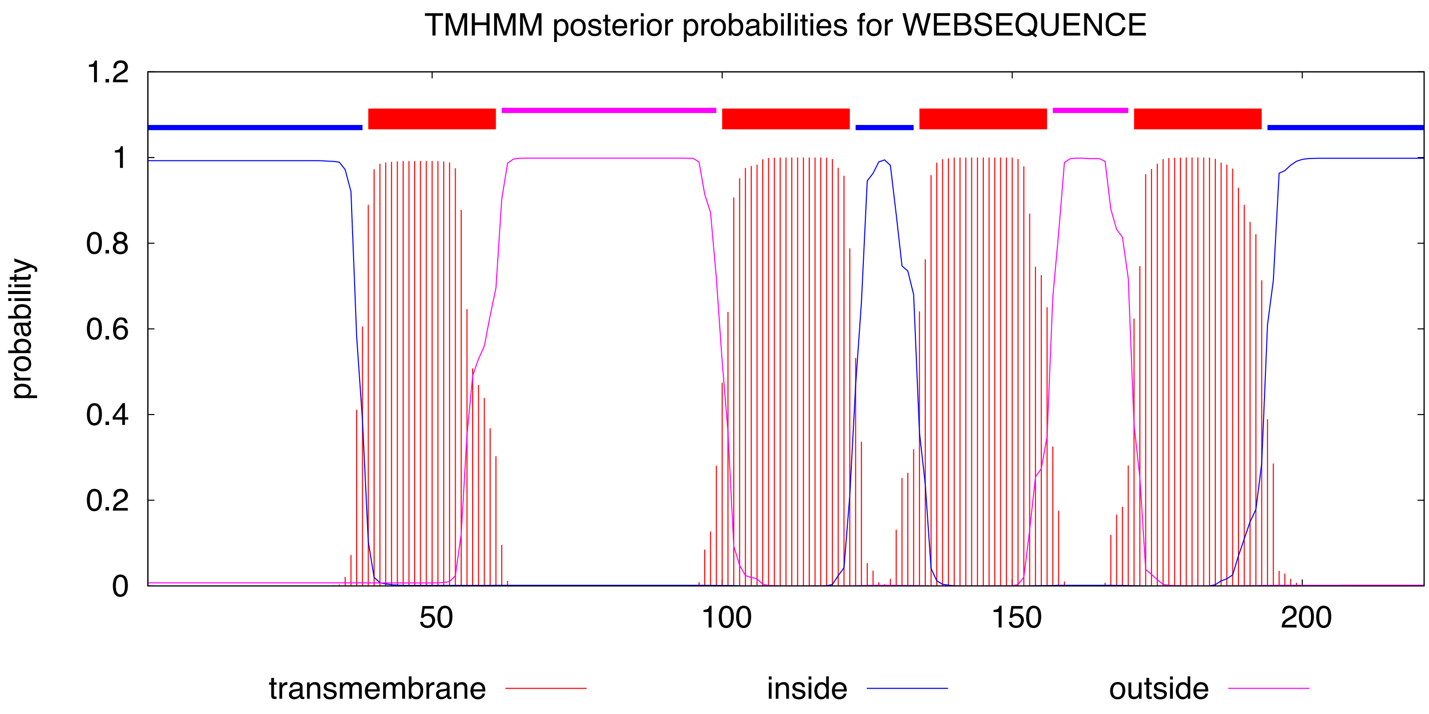

**Supplementary Figure S2**

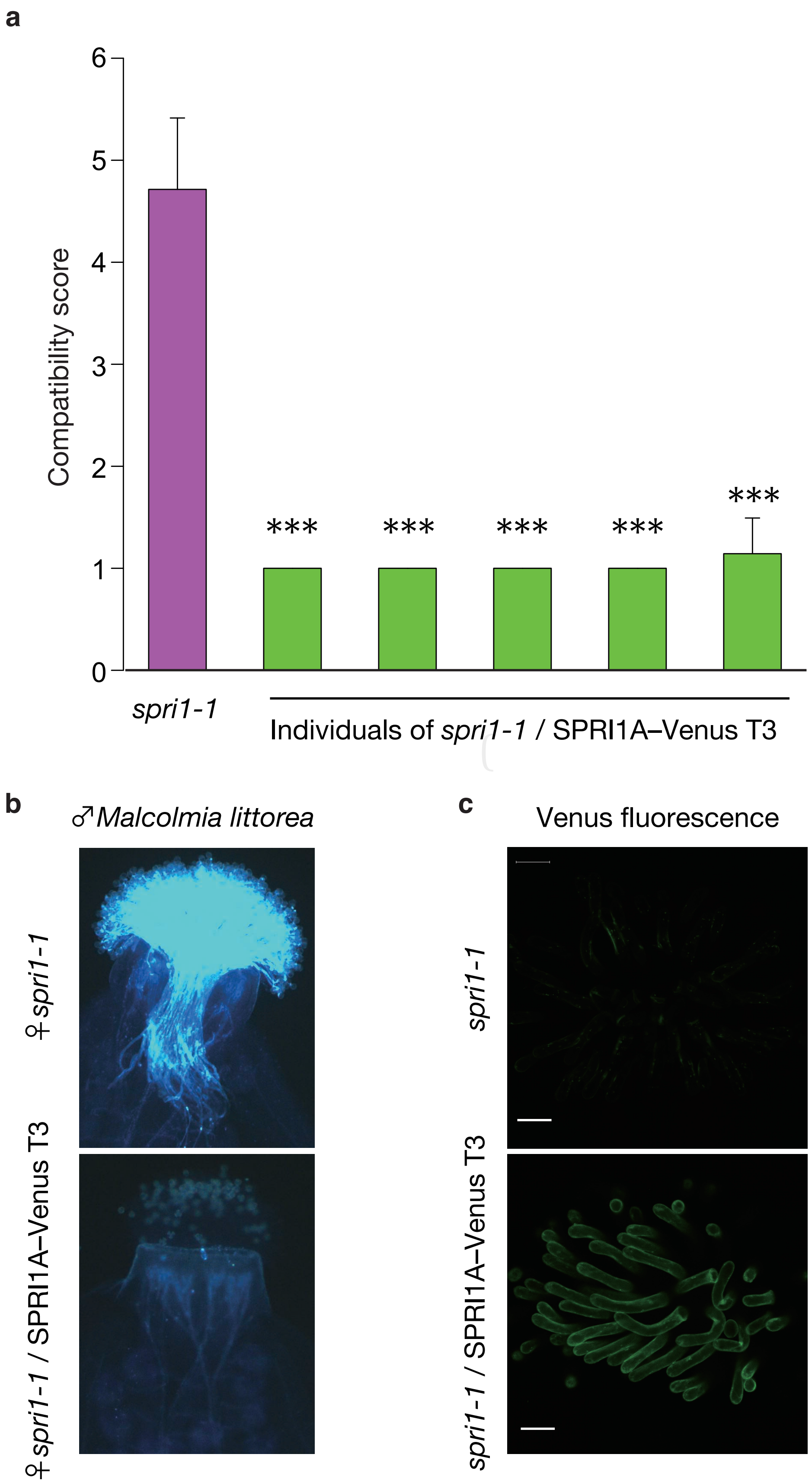

**Supplementary Figure S3**

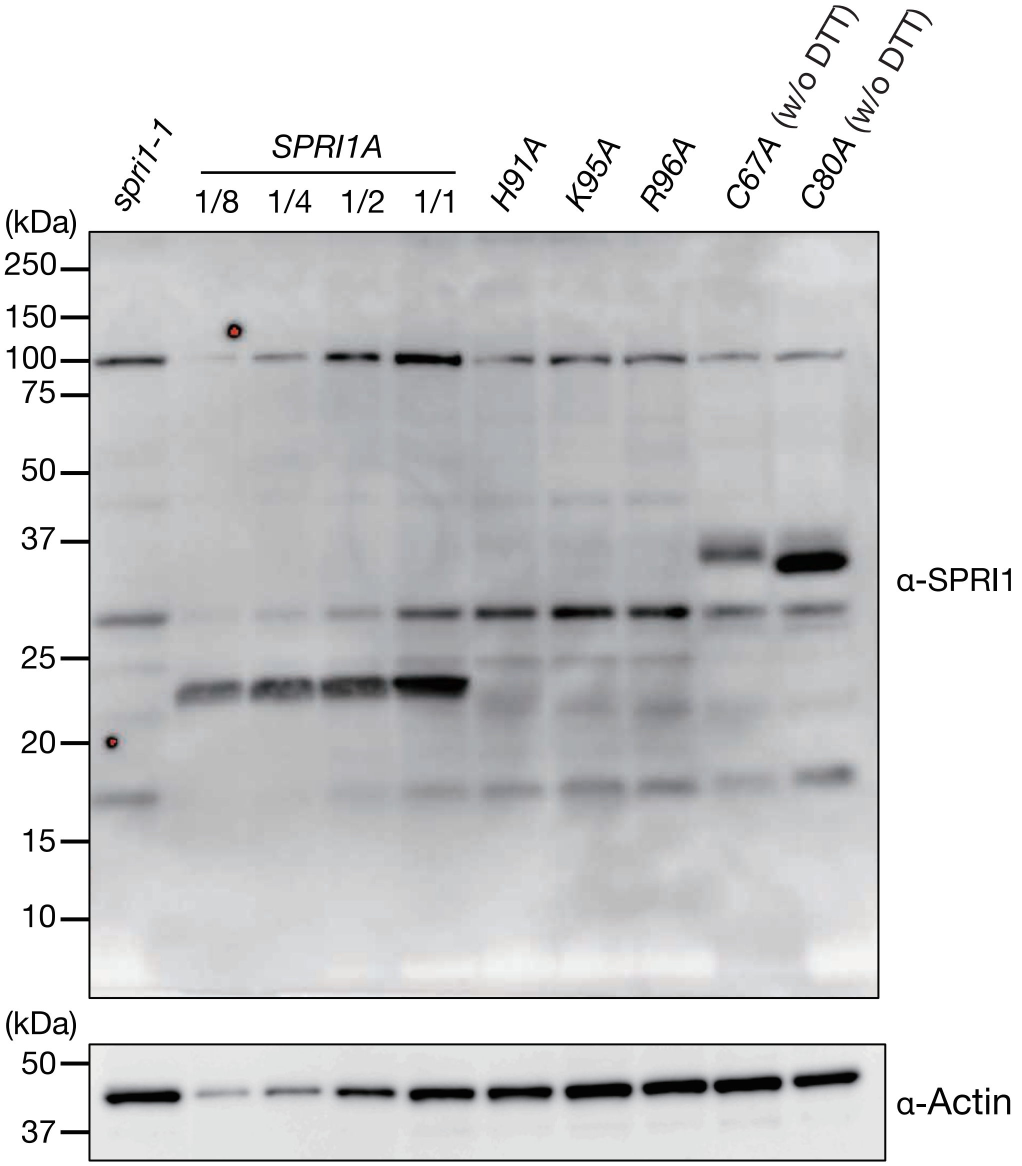

**Supplementary Figure S4**

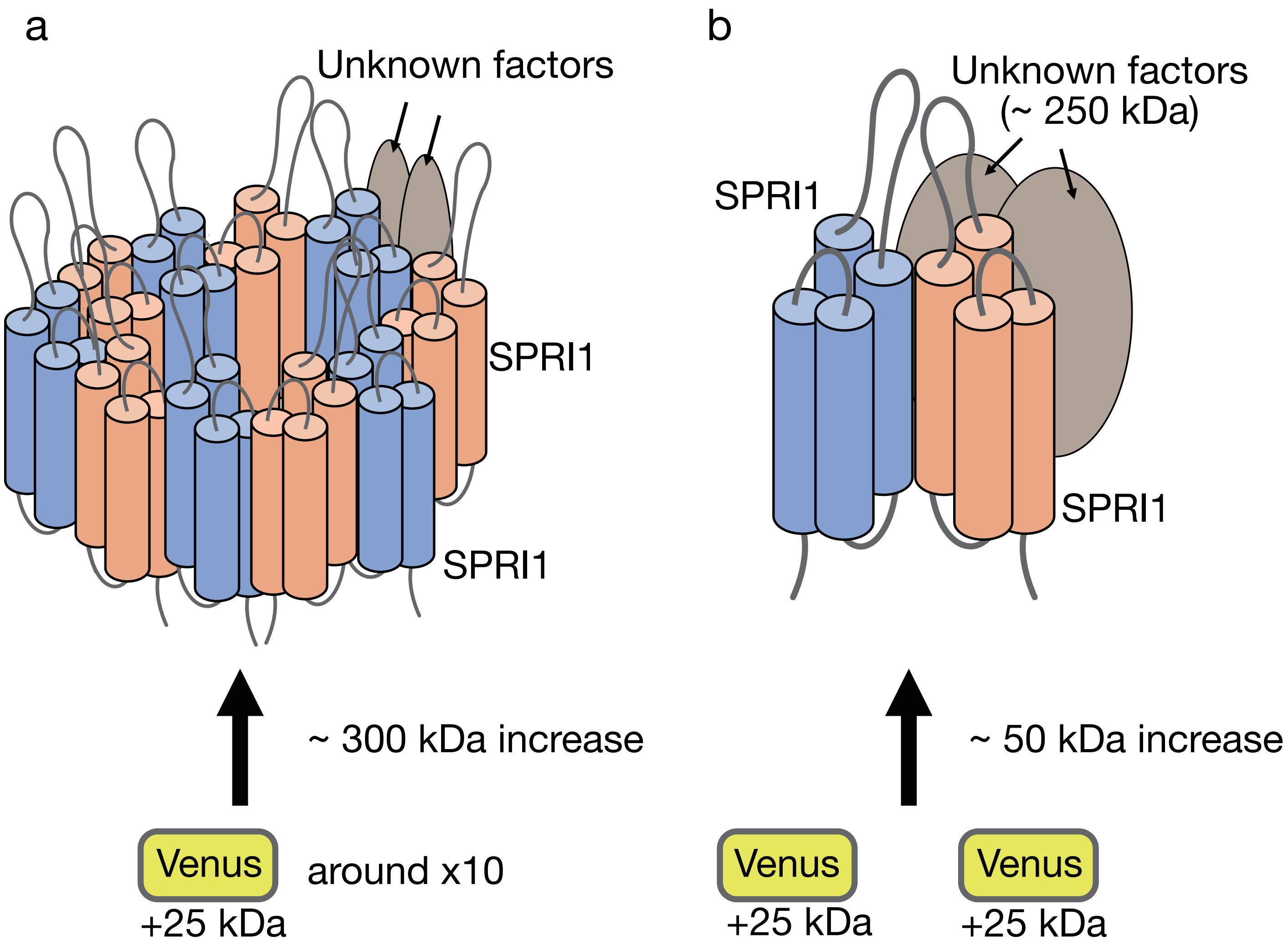

**Supplementary Figure S5**

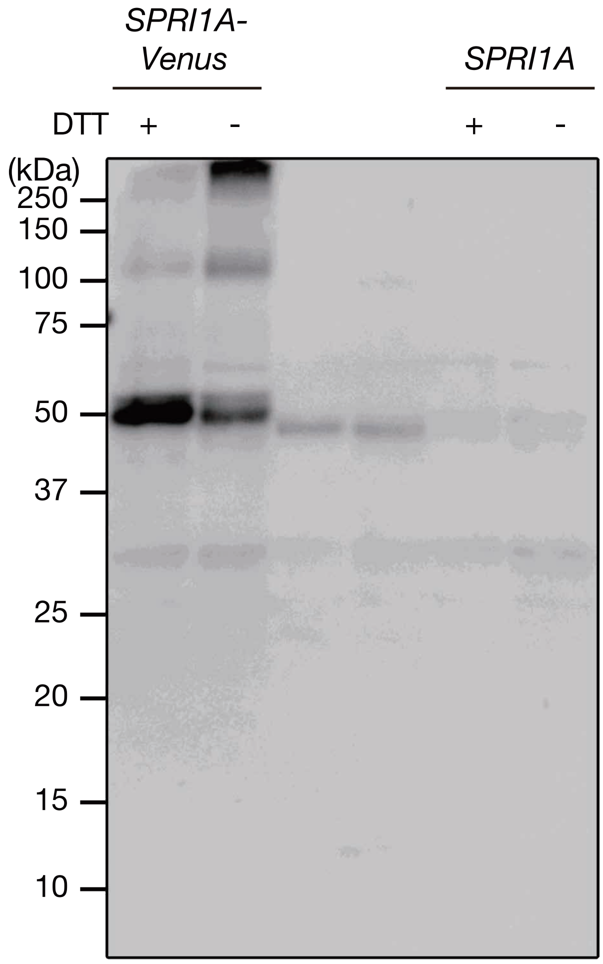
